# Imp/Syp temporal gradients govern decommissioning of *Drosophila* neural stem cells

**DOI:** 10.1101/136655

**Authors:** Ching-Po Yang, Tamsin J. Samuels, Yaling Huang, Lu Yang, David Ish-Horowicz, Ilan Davis, Tzumin Lee

## Abstract

Timing of *Drosophila* neuroblast decommissioning is controlled in a lineage-specific manner. Following a prepupal ecdysone pulse, the ecdysone receptor and mediator complex cause neuroblasts to shrink. Shrinking is followed by nuclear accumulation of Prospero and cell cycle exit. Only mushroom body (MB) neuroblasts escape early pupal termination. Here, we demonstrate that the opposing temporal gradients of Imp and Syp RNA-binding proteins that govern temporal fate also regulate neuroblast decommissioning. The Imp gradient declines slower in MB neuroblasts so they still express Imp when it is absent from others. The presence of Imp in MB neuroblasts prevents decommissioning partly through inhibiting the mediator complex. Moreover, a timely induction of Imp can protect many non-MB neuroblasts from aging. We also show that the increasing Syp gradient permits Prospero accumulation and neuroblast termination. Together our results reveal that progeny temporal fate and progenitor decommissioning are co-regulated in protracted neuronal lineages.

## Introduction

A long-lived neural stem cell shows age-dependent changes in proliferation and progeny cell fate (Li et al., 2013). Such temporal regulations are lineage-specific in *Drosophila* where individual neuroblasts produce unique series of neuronal types and end neurogenesis independently (Chia et al., 2013; Homem et al., 2015; Lee et al., 1999; Lin et al., 2012; Sousa-Nunes, et al., 2010; Truman and Bate, 1988; Yu et al., 2010). Moreover, the neuronal temporal fate (birth order-dependent cell fate) and neuroblast termination is likely to be co-regulated, to produce the required number and types of neurons in a given lineage before the neuroblast stops dividing (Maurange et al., 2008). This possibly involves a lineage-autonomous developmental clock that governs both progeny temporal fate and neuroblast aging in each neuronal lineage.

Temporal gradients of proteins, including IGF-II mRNA-binding protein (Imp), Syncrip (Syp), and Chinmo, govern neuronal temporal fates in diverse neuroblast lineages (Liu et al., 2015; Zhu et al., 2006). Imp and Syp are RNA-binding proteins, and Chinmo is a BTB zinc-finger nuclear protein. The descending Imp and ascending Syp gradients co-exist in neuroblasts that confer serially derived neurons with birth order-dependent opposite levels of Imp and Syp. Imp enhances and Syp represses Chinmo translation, which results in a Chinmo protein gradient among the newborn neurons. High Chinmo promotes early neuronal temporal fates. The opposing Imp/Syp temporal gradients extend throughout neurogenesis but exhibit distinct lineage-characteristic temporal dynamics (Liu et al., 2015). In the antennal lobe (AL) lineages, one neuroblast can yield a sequence of 20+ neuronal types from ~80 asymmetric cell divisions that span over four days of larval development (Yu et al., 2010; Lin et al., 2012). The opposite Imp/Syp temporal gradients are steep and progress rapidly in such fast changing neuroblasts. By contrast, the neuroblasts, which make neurons constituting the mushroom body (MB) learning and memory center, divide incessantly throughout larval and pupal development but produce only three morphological classes of neurons (Lee et al., 1999). Imp and Syp are expressed in shallow, slowly progressing gradients in the long-lived MB neuroblasts. The close correlation between the progression of Imp/Syp gradients and the progeny’s temporal fate changes argues for direct coding of diverse neuronal temporal fates by distinct levels of Imp and/or Syp.

Neuroblast termination is also temporally regulated. During active cycling, neuroblasts regrow promptly following each asymmetric cell division (Knoblich, 2008; San-Juán and Baonza, 2011; Song and Lu, 2011). Upon pupation, all neuroblasts, except the four pairs of MB neruoblasts, undergo reduction in size while cycling due to sluggish regrowth (Homem et al., 2014). The size-reduced neuroblasts divide slowly and eventually divide symmetrically into two post-mitotic cells. The stage-specific progressive termination of neuroblasts is apparently an actively regulated process, which we refer to as ‘decommissioning’. The initiation of neuroblast decommissioning by the prepupal ecdysone pulse requires the ecdysone receptor and core components of the mediator complex. They promote key enzymes to increase oxidative phosphorylation in energy metabolism that slows down neuroblast regrowth (Homem et al., 2014). Subsequent nuclear accumulation of Prospero then drives cell cycle exit of neuroblasts (Maurange et al., 2008). Notably, neighboring neuroblasts have individual schedules to exit cell cycle at specific times around 24h after pupal formation (APF) (Awasaki et al., 2014). Moreover, despite the global ecdysone action, the MB neuroblasts remain active in regrowth and cycle for ~100 more divisions at the pupal stage (Ito and Hotta, 1992; Truman and Bate, 1988). The MB neuroblasts finally shrink but terminate via apoptosis around 96h APF (Siegrist et al., 2010). The mechanisms underlying such temporal differences in neuroblast decommissioning remain unexplored.

Given the necessity for producing a whole series of neuronal types from each neuronal lineage, neuroblast decommissioning may be temporally regulated by the mechanisms that govern neuronal temporal fate progression. Here we examined if and how the opposite Imp and Syp temporal gradients, which control neuronal temporal cell fates across protracted neurogenesis, also guide the course of neuroblast decommissioning. We found that Imp governs neuroblasts’ readiness for decommissioning in early pupae, and that Syp enables Pros accumulation in shrunk neuroblasts to end the cell cycle. The MB neuroblasts are insensitive to the prepupal ecdysone pulse due to a protracted gradient of Imp that prevents decommissioning partly through inhibiting the mediator complex. Other neuroblasts lacking Imp shrink rapidly in early pupae, and exit the cell cycle upon Syp-dependent nuclear accumulation of Pros. Imp is dominant to Syp, as high Syp promotes Pros-dependent cell cycle exit only in those Imp-lacking decommissioned neuroblasts. Taken together, neuroblast decommissioning is an actively programmed developmental process that is temporally regulated by the opposite Imp/Syp gradients in each neuroblast to ensure its completion of an entire lineage.

## Results

### Non-MB NBs undergo early pupal decommissioning due to lack of Imp and exit cell cycles due to high Syp

In order to monitor decommissioning of central brain neuroblasts, we expressed a cell cycle-sensitive GFP using a GAL4 driver that is active in all, but not optic lobe, neuroblasts. In the wild-type control, almost all neuroblasts had disappeared by 48 hours APF and only four neuroblasts per hemisphere remained. These four were the MB neuroblasts, which have been described to divide until 96 hours APF (Siegrist et al., 2010). The MB neuroblasts too were gone by adult stage (Fig. 1A). Leaving the MB NBs out of our analysis, we followed the decommissioning of the remaining neuroblasts (~100 per hemisphere) from wandering larvae to early pupae. The average size had decreased significantly by 8 hours APF and most neuroblasts had reduced to a size comparable to that of their daughter ganglion mother cells (GMC) within 24 hours APF (Fig. 1A,E). Notably, the aged neuroblasts did not exit the cell cycle in a synchronized wave. A few neuroblasts (though small in size) remained at 24 hours APF (Fig. 1A’’), as evidenced by the expression of Miranda (Mira), a neuroblast-specific gene, and the occasional presence of pH3, indicating mitosis.

**Fig.1.**
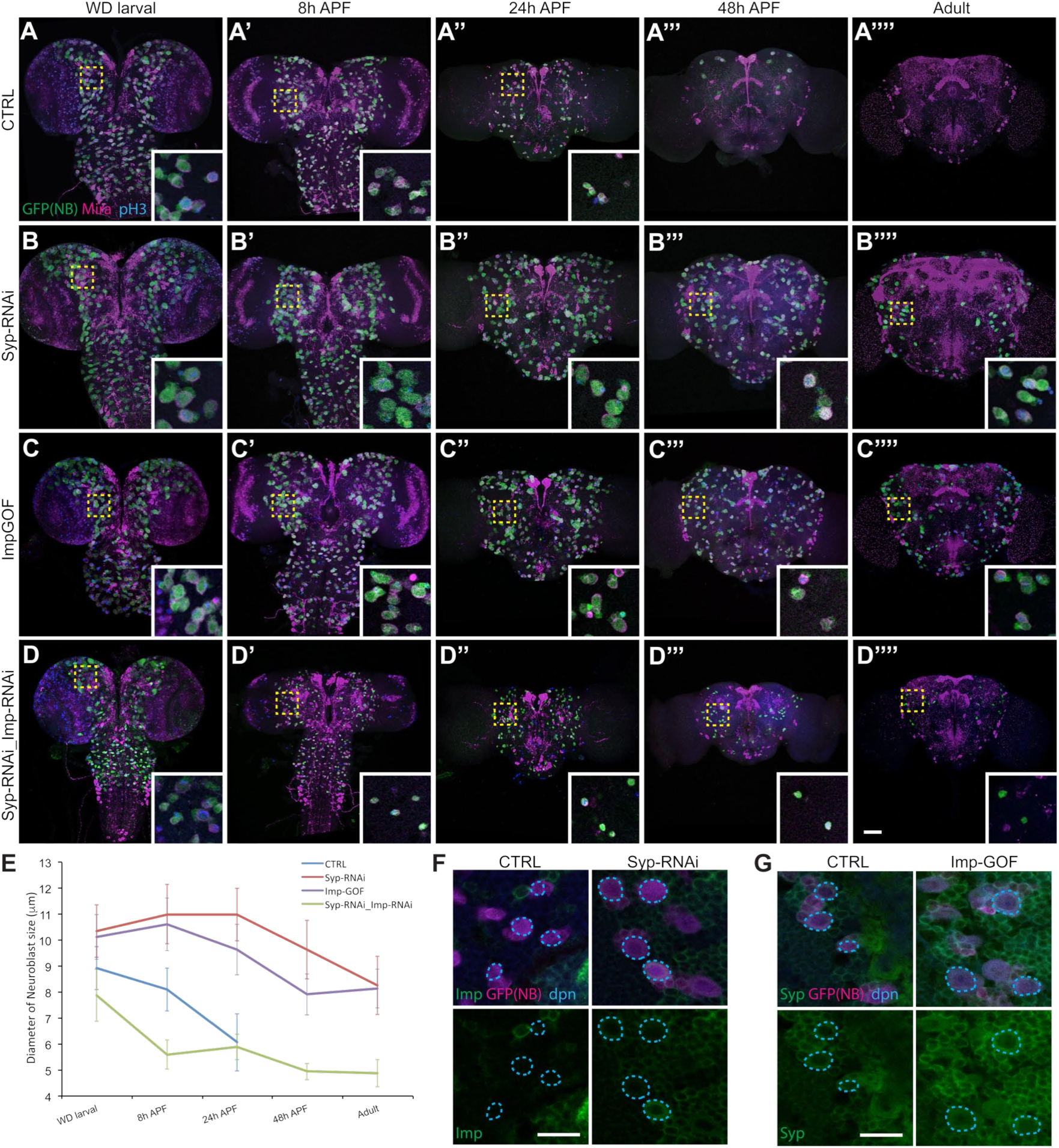
Imp and Syp regulate non-MB NB decommissioning. (A-D) Ectopic Imp or Syp-depleted prolong NB life into adult. Composite confocal images of CTRL (A-A’’’’), Syp-depleted (B-B’’’’), Imp gain-of-function (C-C’’’’), and Syp/Imp-depleted fly brain at specific developmental ages were counterstained with GFP (Green), Mira (Magenta) and phospho Histone H3 (pH3, Blue). Transgenes were driven by *dpnEE-GAL4* in the neuroblasts of central brain. Scale bar: 50μm. (E) Quantification of the size of neuroblast in the anterior region of fly brain (measured by the diameter of Mira labeled neuroblasts, mean ± SD, n=6 brains) from (A-D). (F) Prolonged Imp expression in Syp-depleted neuroblasts. Representative confocal images of 8h APF fly brain counterstained with Imp (Green), GFP (magenta) and dpn (Blue) in CTRL and Syp depletion flies. Scale bar: 20μm. (G)Imp overexpression did not affect Syp expression. Representative confocal images of 8h APF fly brain counterstained with Syp (Green), GFP (magenta) and dpn (Blue) in CTRL and Imp gain-of-function flies. Scale bar: 20μm.

Those decommissioned neuroblasts in early pupae were negative for Imp and positive for Syp. We wondered if neuroblasts purposely locked in their beginning state of Imp/Syp expression could escape decommissioning. We tested this by silencing Syp with targeted RNAi, which secondarily maintained high Imp throughout neuroblast life (Fig. 1F). We found that high Imp and minimal Syp drastically delayed neuroblast decommissioning (Fig. 1B). Most, if not all, late larval neuroblasts remained at 48h APF; quite a few sustained and continued to cycle at the adult stage (Fig. 1B’’’’). The size of Syp-depleted neuroblasts was not reduced by 24hours APF, and those who persisted were consistently larger than GMCs (Fig. 1E). Expressing transgenic Imp continuously elicited similar phenotypes (Fig. 1C). Notably, Syp existed abundantly in those Imp-overexpressing neuroblasts that showed no evidence of aging in early pupae (Fig. 1G). This argues that it is ectopic Imp, rather than absence of Syp, which accounts for the suppression of early pupal neuroblast decommissioning in both loss-of-Syp and gain-of-Imp conditions.

Consistent with Imp dominantly repressing neuroblast decommissioning, silencing Imp on top of Syp RNAi restored the early-pupal neuroblast decommissioning (Fig. 1D). Neuroblasts with depleted Imp and Syp underwent accelerated shrinkage in early pupae, indicating a prompt aging in response to the prepupal ecdysone pulse (Fig. 1E).

However, the aged neuroblasts failed to terminate until late pupal or even adult stage (Fig. 1D’’’’). Together, we hypothesize that Imp levels determine if neuroblasts shrink in early pupae. Once decreased in size, the neuroblasts require high Syp to exit the cell cycle.

### MB NBs escape early pupal decommissioning due to protracted Imp expression

At the late larval stage, neuroblasts no longer express Imp, with the exception of the MB neuroblasts, which maintain detectable Imp around pupation (Fig. 2A). We therefore tested whether Imp expression in the MB neuroblasts is responsible for their long life. Indeed, targeted RNAi rendered Imp undetectable in larval MB neuroblasts and resulted in the termination of MB neurogenesis promptly in early pupae (Fig. 2 B,C and data not shown). Without Imp, the MB neuroblasts were relatively small but stable in size until pupation when they rapidly shrank (Fig. 4K). The MB neuroblasts with a drastically reduced cell size were never found positive for pH3, but a majority of them survived beyond 48 hours APF.

**Fig.2.**
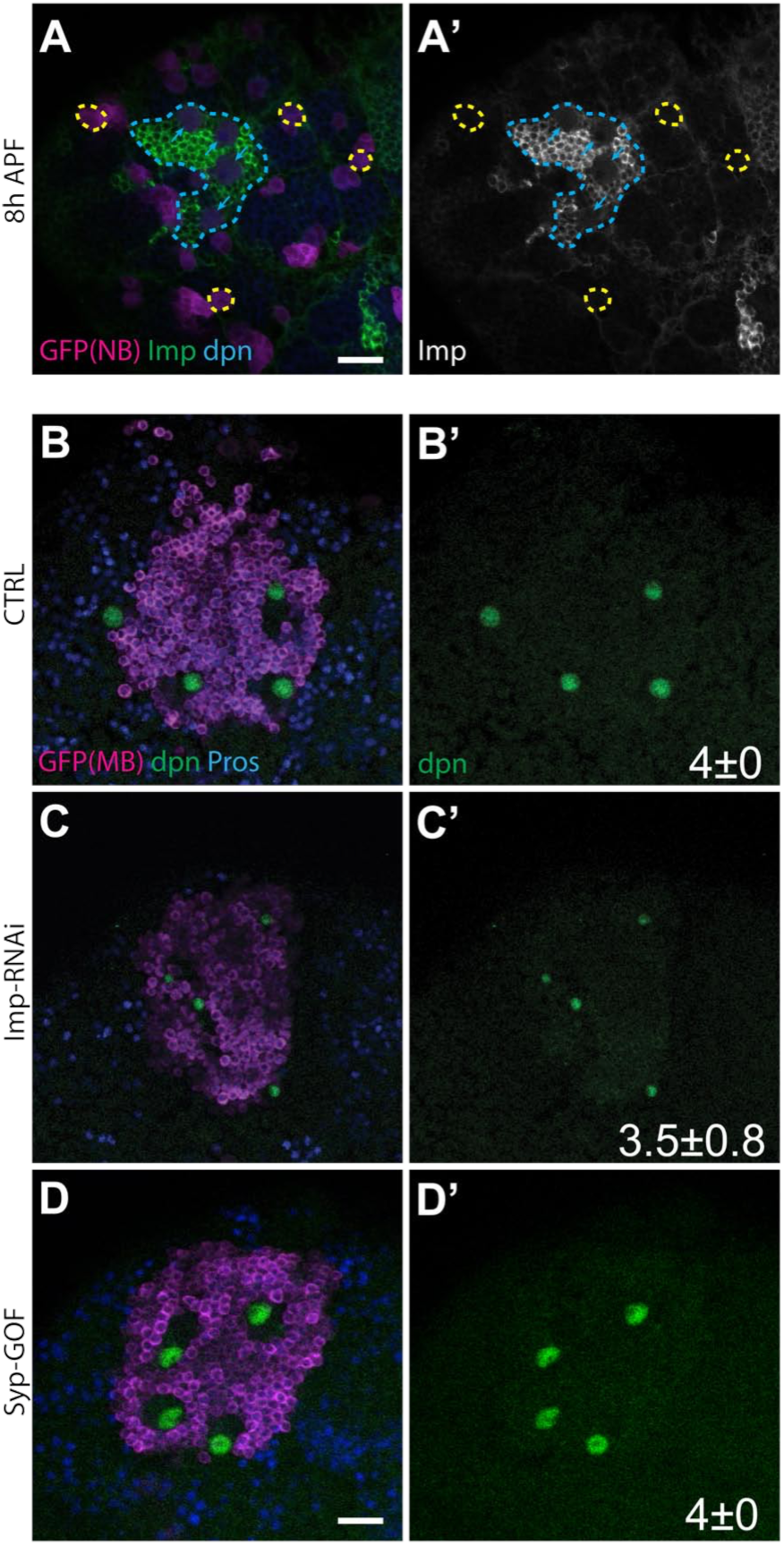
Protracted Imp expression protects MB NBs from early pupal decommissioning. (A) Imp continuously expressed in MB NBs at early pupal development. Representative confocal images of 8h APF fly brain counterstained with Imp (Green), GFP (magenta) and dpn (Blue) in wild-type flies. The blue dots circle the MB region, and arrows indicate protracted Imp expression in the MB NBs. The yellow dots circle the NBs at the same focal plan in the posterior region, which are negative for Imp expression. Scale bar: 20μm. (B-D) Imp depletion prematurely ended MB neurogenesis. Representative confocal images of 48h APF fly brains counterstained with dpn (Green), GFP (magenta) and Pros (Blue) in CTRL (B), Imp depletion (C), and Syp gain-of-function (D). Transgenes were driven by *GAL4-OK107*; numbers of MB NB per brain lobe (mean ± SD, n=6) are indicated at the bottom right of each panel. Scale bar: 20μm.

Wild-type MB neuroblasts also express a much lower level of Syp than non-MB neuroblasts (Liu et al., 2015 and data not shown). However, overexpressing Syp in MB neuroblasts could not cause these cells to undergo decommissioning in early pupae, like other neuroblasts. They outlived non-MB neuroblasts and persisted in normal size into late pupae (Fig. 2D). Taken together, our results implicate prolonged Imp expression, rather than slowly uprising Syp, in protecting wild-type MB neuroblasts from early pupal decommissioning.

### Late-larval induction of transgenic Imp effectively protects non-MB NBs from early pupal decommissioning

It is possible that MB and non-MB neuroblasts undergo progressive aging at different speeds during larval development and only the non-MB neuroblasts are aged enough to undergo decommissioning in response to the prepupal ecdysone pulse. If true, overexpressing Imp since the beginning of neurogenesis could slow down the continuous aging of neuroblasts and indirectly affect neuroblasts’ responses to ecdysone. Alternatively, Imp may directly regulate the ecdysone signaling and/or its downstream effectors. To address this, we tried not to perturb the normal larval aging (if any) of neuroblasts and examined if late larval induction of transgenic Imp could protect non-MB neuroblasts from early pupal decommissioning.

We determined if there is a critical window for transgenic Imp to sustain non-MB neuroblasts into late pupae. We controlled the timing of transgenic Imp expression in all neuroblasts positive for dpnEE-GAL4 (Awasaki et al., 2014), using a temperature-sensitive GAL80 (McGuire, et al., 2003). Transgenic Imp was induced at a specific time around pupation with a temperature shift from 18°C to 29°C (Fig. 3A). Induction at 8h before pupa formation (BPF) or later failed to extend any non-MB neuroblasts beyond early pupal development (Fig. 3C and data not shown). Induction at 12h BPF, by contrast, substantially prolonged the life of a few non-MB neuroblasts (Fig. 3D); and many more neuroblasts could survive into late pupae following induction of Imp at 15h BPF (Fig. 3E). Induction earlier than 15h BPF did not further increase the number of neuroblasts persisting in mid-pupal brains (Fig. 3F,G). We could not rescue most non-MB neuroblasts as in the case with continuous Imp expression, probably due to a mosaic inactivation of GAL80[ts] as evidenced by immunostaining of Imp (data not shown). The potency of late larval Imp induction in sustaining many non-MB neuroblasts suggests that Imp directly suppresses the activation of the early pupal neuroblast decommissioning by the prepupal ecdysone pulse.

**Fig.3.**
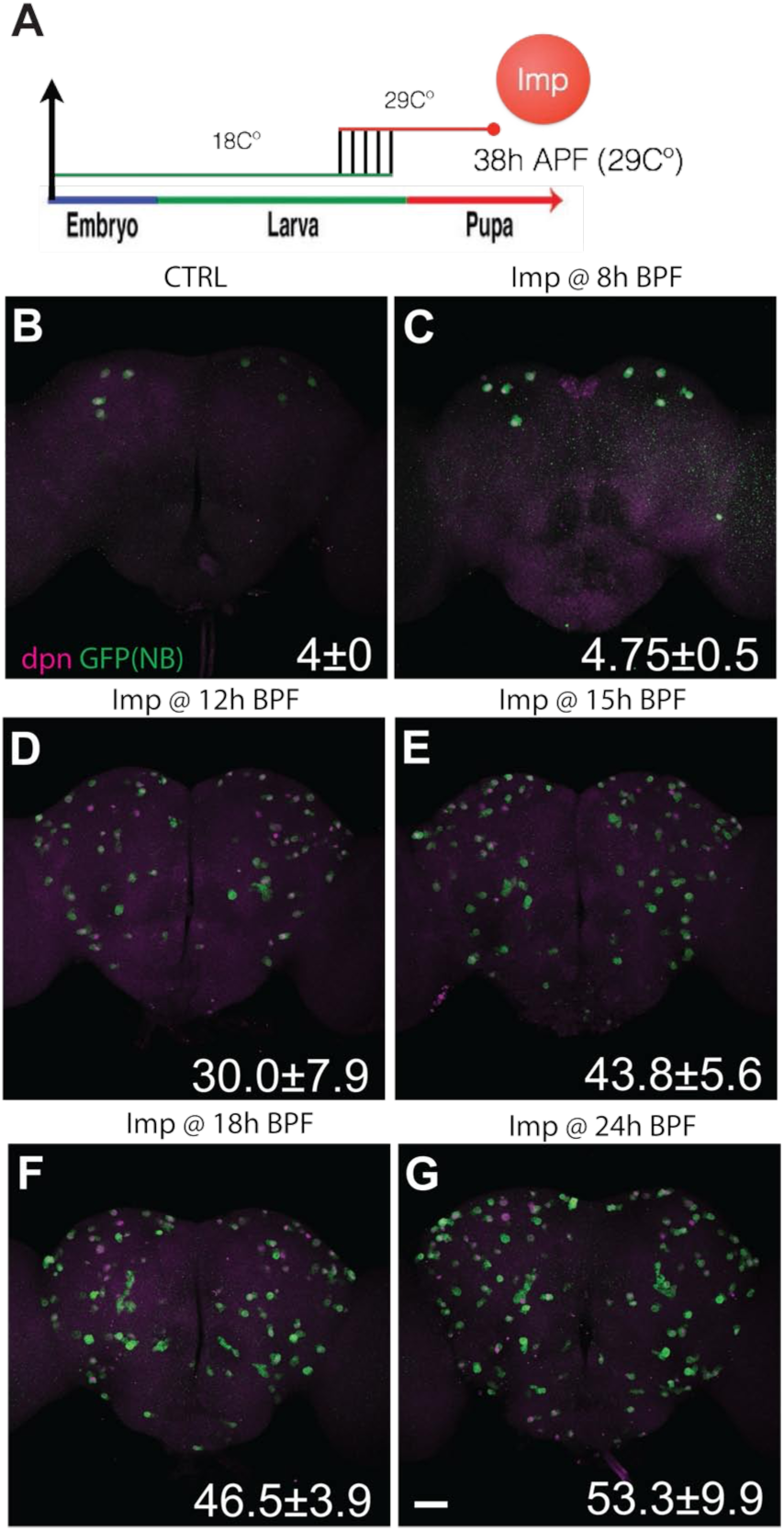
Induction of transgenic Imp at late larval stage prolongs NB life. (A) Scheme for targeted induction of transgenic Imp by heat shock to inactive the temperature-sensitive GAL80 at specific developmental times. Flies were continuously incubated at 29°C after heat shock and fly brains were dissected and fixed after 38hr pupal formation (38h APF). (B-G) Late larval induction of Imp prolongs NB life. Composite confocal images of CTRL (B) and targeted transgenic Imp induction at 8h (C), 12h (D), 15h (E), 18h (F) and 24h (G) BPF fly brains were counterstained with GFP (Green) and dpn (Magenta). Number of NB per brain lobe (mean ± SD, n=6) are indicated at the bottom right of each panel. Scale bar: 50μm.

**Fig.4.**
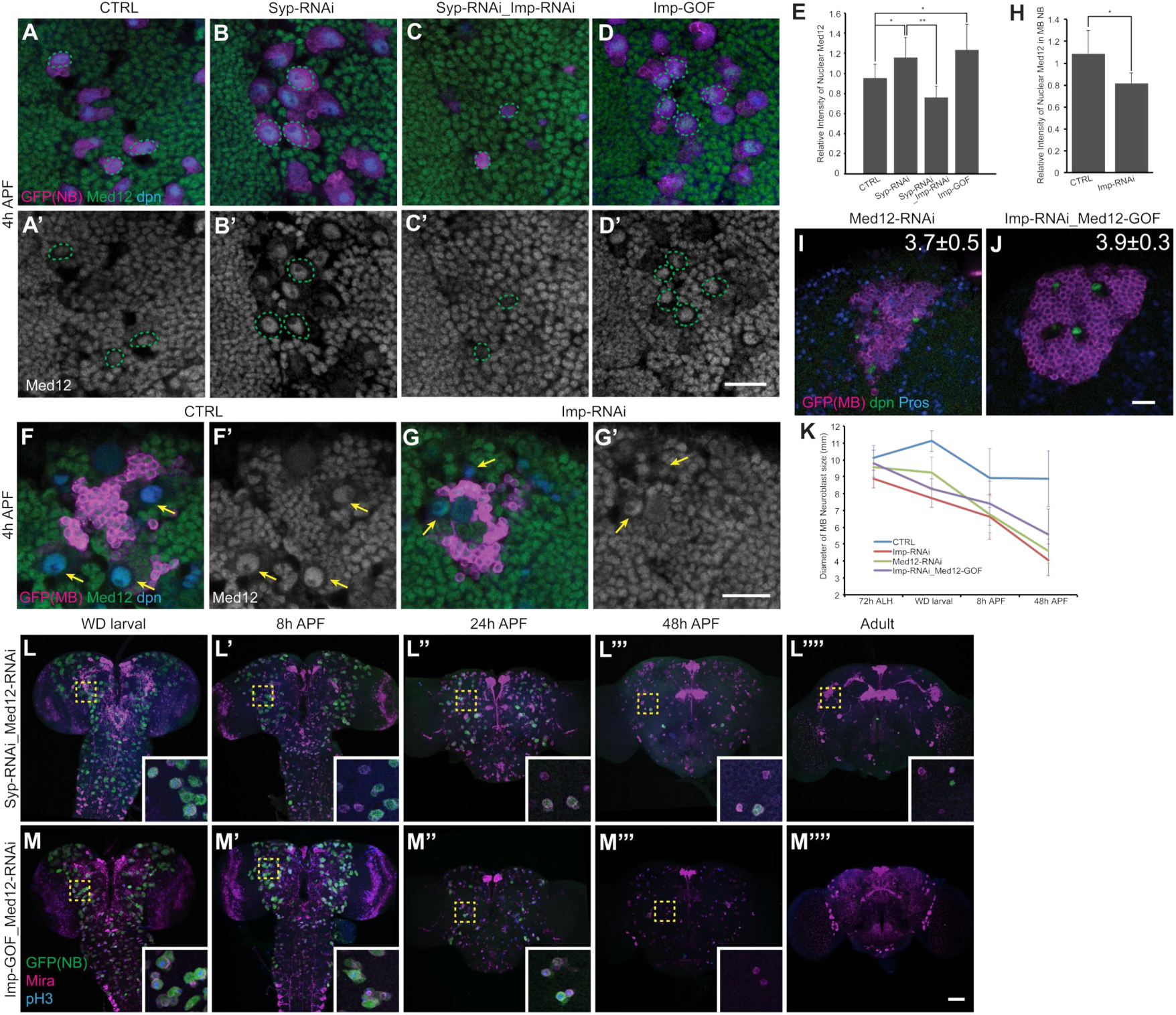
Imp suppresses NBs shrinkage through inhibiting mediator complex. (A-D) Syp suppresses Med12 expression in NB. Representative confocal images of 4h APF fly brain counterstained with Med12 (Green), GFP (magenta) and dpn (Blue) in CTRL (A), Syp depletion (B), Syp/Imp depletion (C) and Imp gain-of-function (D). Green dots circle the nuclear staining of Med12 in the neuroblast. Scale bar: 20μm. (E) Quantification of the relative intensity of Med12 in the anterior region of fly brain from (A-D), two-tailed Student’s t test was performed. *p< 0.05, **p< 0.01. Relative Med12 intensity was determined by the averaging signals in the neuroblast followed by normalization to the averaging signals in the optic lobe region (mean ± SD, n=6 brains). (F,G) Imp promotes Med12 expression in MB NB. Representative confocal images of 4h APF fly brains counterstained with Med12 (Green), GFP (magenta) and dpn (Blue) in CTRL (E) and Imp depletion (F). Arrows indicated the nuclear staining of Med12 in the MB neuroblast. Scale bar: 20μm. (H) Quantification of the relative intensity of Med12 in the MB NBs from (F) and (G), two-tailed Student’s t test was performed. *p< 0.05 (mean ± SD, n=6 brains). (I-K) Mediator complex is downstream of Imp to regulate MB NBs shrinkage. Representative confocal images of 48h APF fly brains counterstained with dpn (Green), GFP (magenta) and Pros (Blue) in Med12 depletion (I) and Imp-depleted, Med12 gain-of-function (J) flies. Numbers of MB NB per brain lobe (mean ± SD, n=6) are indicated at the top right of each panel. Scale bar: 20μm. (K) Quantification of the size of MB neuroblast (measured by the diameter of Mira signal (mean ± SD, n=6 brains)) from other samples preparation of transgenes driven by *GAL4-OK107*. (L,M) Mediator complex is downstream of Imp to induce NBs shrinkage. Composite confocal images of Syp/Med12 knockdown (L) and Imp gain-of-function, Med12 knockdown (M) fly brain at specific developmental ages counterstained with GFP (Green), Mira (Magenta) and phospho Histone H3 (pH3, Blue). Scale bar: 50μm.

### Imp suppresses NB decommissioning partly through inhibiting mediator complex

The ecdysone receptor and mediator complex have been recently implicated in NB shrinking and ultimately ending (Homem et al., 2014). The mediator complex is essential for an energy metabolism switch that slows down neuroblast regrowth during successive divisions and thus progressively reduces the neuroblast cell size. The NB shrinkage and termination could also be delayed by expressing the inhibition subunit of mediator complex, Med12 (Kohtalo, Kto) (Homem et al., 2014; Malik and Roeder, 2010).

To examine the possibility that Imp may prevent neuroblast shrinkage via suppressing the mediator complex, we determined Med12 expression in the neuroblasts following various Imp/Syp manipulations. We found elevated Med12 protein levels in non-MB neuroblasts with Syp RNAi around pupation (Fig. 4A,B,E). The Med12 protein levels were reduced to lower than normal upon silencing Imp on top of Syp RNAi (Fig. 4C,E). The Imp-depleted MB neuroblasts also showed a significant reduction in Med12 expression (Fig. 4F-H). In sum, the inverse correlation between Med12 protein levels and early pupal neuroblast shrinkage was seen in all Imp/Syp manipulations.

In further support of the mediator complex acting downstream of Imp/Syp, silencing Med12 restored shrinkage of Syp-depleted non-MB neuroblasts in early pupae (Fig. 4L). However, the size-reduced neuroblasts (positive for Mira despite a gradual loss of the neuroblast labeling by dpnEE-GAL4) remained defective in cell cycle exit and sustained with evidence of cycling even at adult stage (Fig. 4L’’’’), reminiscent of the perpetuation of those tiny non-MB neuroblasts caused by repression of both Imp and Syp (Fig. 1D’’’’). By contrast, silencing Med12 could make Imp-overexpressing neuroblasts not only shrink but also end cell cycles by 48h APF (Fig. 4M). Silencing Med12 alone was also sufficient to make MB neuroblasts shrink rapidly in early pupae, similar to non-MB neuroblasts (Fig. 4I). These results suggest that Imp suppresses neuroblast decommissioning via inhibiting the mediator complex.

Despite the notion that Imp increases Med12 expression and secondarily prevents neuroblasts shrinkage (Fig. 4D,E), overexpressing Med12 only partially blocked the rapid shrinkage of Imp-depleted MB neuroblasts in early pupae (Fig. 4J). This implies that Imp may suppress the mediator complex via not only enhancing inhibitors (e.g. Med12) but also repressing activators (e.g. various core components of the mediator complex).

Eight mediator complex components, including Med4, Med6, Med9, Med10, Med11, Med22, Med27, and Med31, have been implicated in promoting neuroblast shrinkage following pupation (Homem et al., 2014). In order to determine whether *Med12* and other *Med* transcripts are targets of the Imp RNA-binding protein, we pulled down endogenous Imp::GFP (Toledano, et al., 2012) and measured transcript levels using RIP-qPCR. We found that the *Med6* mRNA was enriched to the largest extent (Fig. 5A). However, silencing Med6 did not show stronger phenotypes than manipulating other Mediator genes (Fig. 5B, and data not shown). By contrast, the *Med12* mRNA was not significantly enriched compared to the negative control *rp49* (Fig. 5A). In all cases the control GFP immunoprecipitation from a wild type strain was very clean, ranging from 0.09% to 0.54% of input (data not shown). Unlike Imp overexpression that could sustain many neuroblasts into adults, compromising the mediator complex alone failed to extend neuroblasts beyond the mid-pupal stage (Fig. 5B). Collectively, these results suggest that Imp prevents neuroblast decommissioning only partly through inhibiting the mediator complex, and that it is also likely to involve additional unknown components.

**Fig.5.**
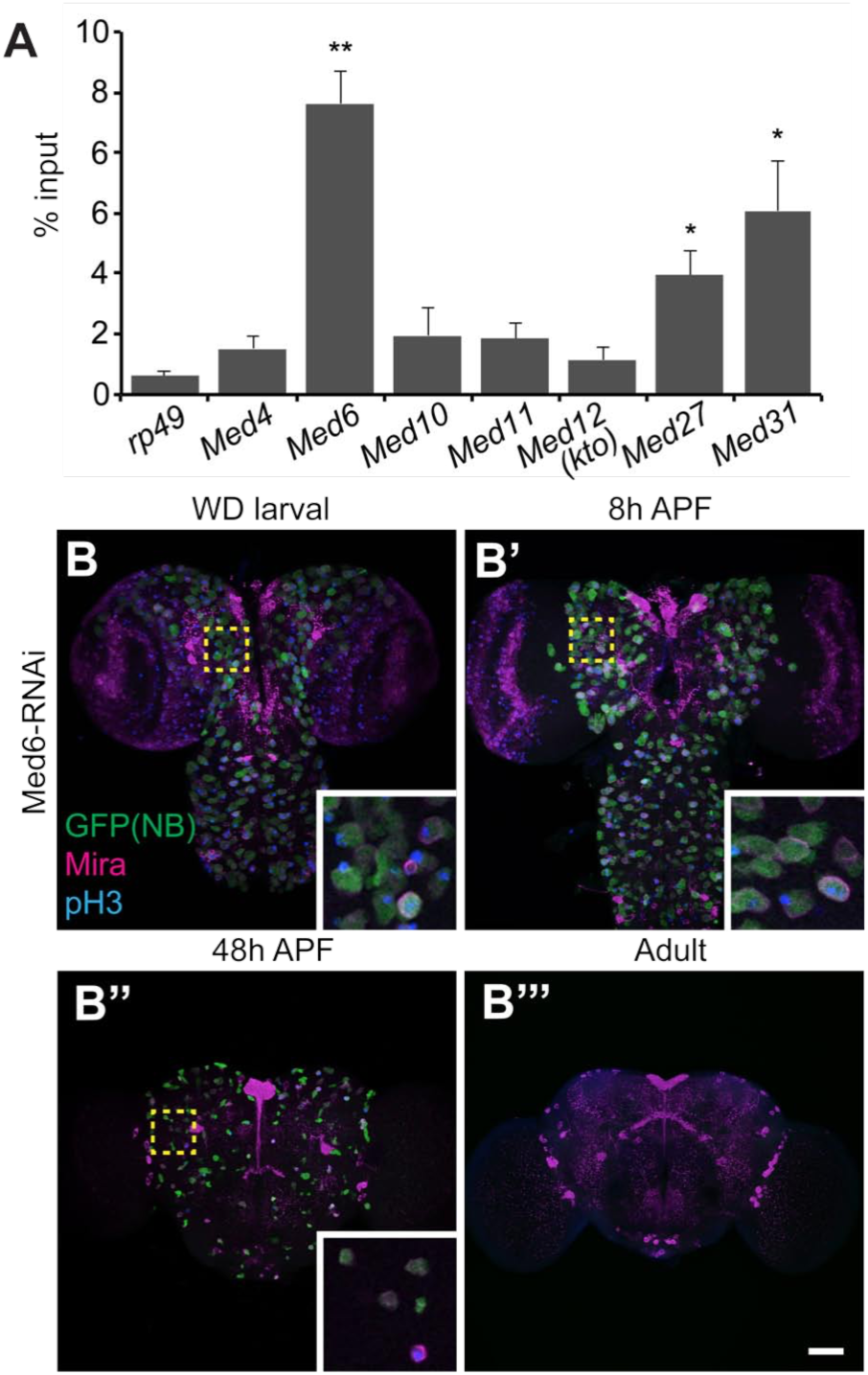
Med6 as a potential direct target of Imp. (A) Med6, but not Med12, mRNA could be effectively pulled down with Imp. RT-qPCR (mean ± s.e.m., n=3; the “x” indicated the individual data points) of RNA immunoprecipitation from Imp::GFP larval brain lysates shows the percent inputs of candidate mediator complex components that were recovered by anti-GFP immunoprecipitation, but not the Med12. Two-tailed Student’s t test was performed comparing the recovery of each transcript to that of the non-binding *rp49*. *p< 0.05, **p< 0.01. (B) Silencing Med6 prolongs NB life. Composite confocal images of Med6-depleted fly brain at specific developmental ages were counterstained with GFP (Green), Mira (Magenta) and phospho Histone H3 (pH3, Blue). Scale bar: 50μm.

### High Syp promotes NB cell cycle exit by permitting nuclear Pros accumulation

Aged neuroblasts with reduced cell size further require Syp to exit cell cycle (Figs, 1D,4L). In wild type, nuclear accumulation of Pros precedes and promotes neuroblast cell cycle exit (Maurange et al., 2008). Moreover, Syp protein may bind directly with Pros mRNA as evidenced by great enrichment of Pros mRNA in Syp IP (McDermott et al., 2014). These phenomena raised the possibility that Syp RNAi delayed neuroblast cell cycle exit via preventing Pros accumulation. Locating Pros proteins in wild-type larval/pupal brains revealed a drastic increase of Pros among the progenies of late larval or early pupal neuroblasts (Fig. 6A,D). Knocking down Syp by targeted RNAi in neuroblasts suppressed the enhancement of Pros in the late-born progeny (Fig. 6B). Syp RNAi also significantly reduced the modest levels of Pros proteins in late neuroblasts and consistently excluded Pros from entering the nuclei (Fig. 6B,D). It has been shown that acute induction of transgenic Pros can promptly terminate actively cycling neuroblasts (Li and Vaessin, 2000). We found transgenic Pros is equally potent in the termination of otherwise long-lasting Syp-depleted neuroblasts (Fig. 6E-G). These results indicate that high Syp enhances Pros, which in turn promotes neuroblast cell cycle exit.

**Fig.6.**
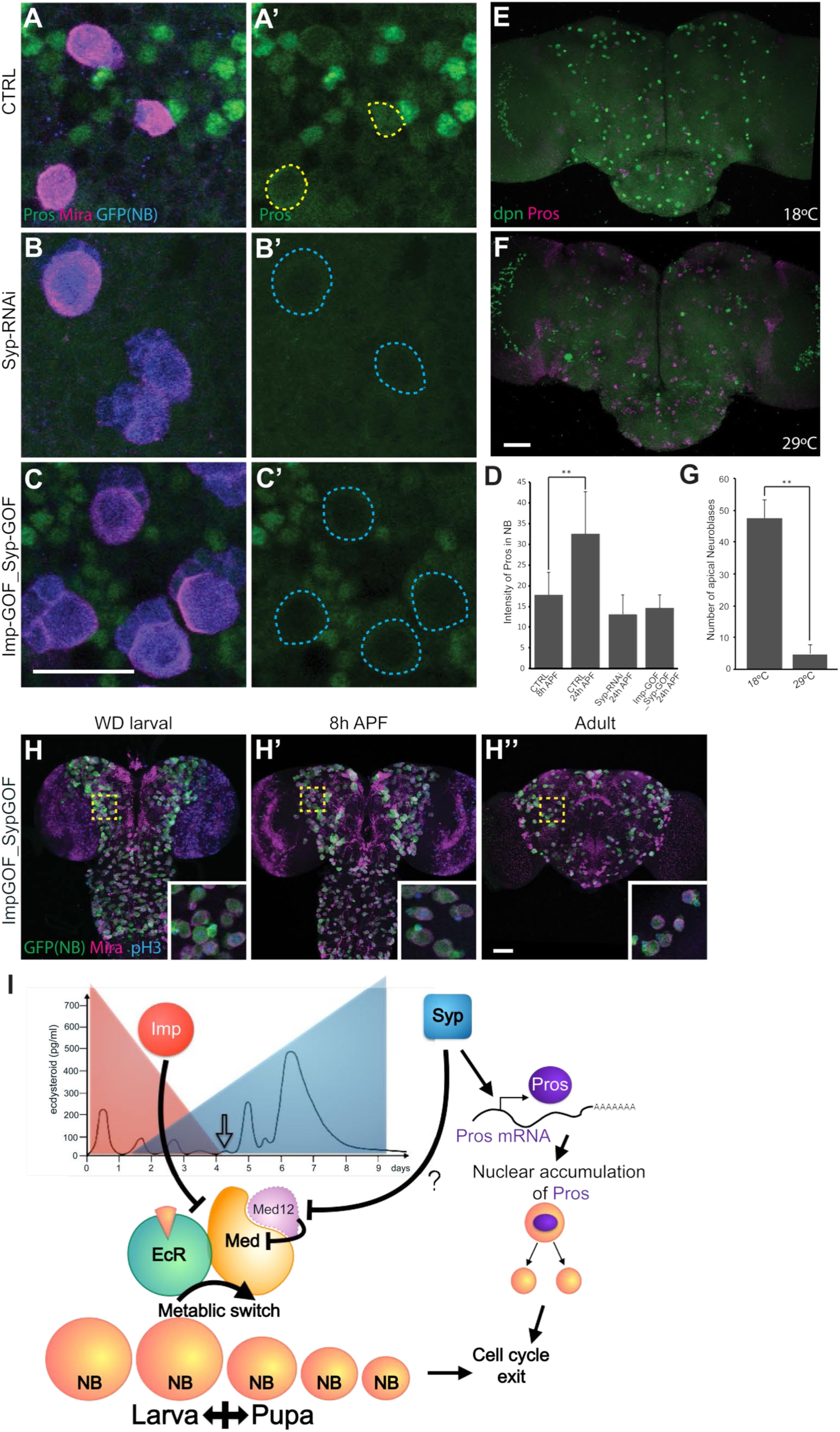
Syp ensures NBs exit by promoting Pros accumulation. (A-C) Syp is required for the accumulation of Pros. Representative confocal images of 24h APF fly brains counterstained with Pros (Green), Mira (magenta) and GFP (Blue) in CTRL (A), Syp depletion (B), and Syp/Imp gain-of-function (C). The neuroblasts were outlined in dashed lines. Scale bar: 20μm. (D) Quantification of the normalized intensity of Pros in the NB from (A-C), two-tailed Student’s t test was performed. **p< 0.01. Pros intensity was determined by the averaging signals in the neuroblast followed by normalization to the averaging signals in the optic lobe region (mean ± SD, n=6 brains). (E,F) Induce Pros misexpression terminated the long-lasting Syp-depleted neuroblasts. Composite confocal images of Syp depletion flies without (E) or with (F) the ectopic induction of Pros at early pupal development were counterstained with dpn (Green) and Pros (Magenta). Scale bar: 50μm. (G) Quantification of the number of neuroblast in the fly brain (measured by the dpn signals (mean ± SD, n=6)) from (E) and (F). Two-tailed Student’s t test was performed. **p< 0.01. (H) Ectopic Imp is dominant to Syp for NB decommissioning. Composite confocal images of Syp/Imp gain-of-function fly brain at specific developmental ages were counterstained with GFP (Green), Mira (Magenta) and phospho Histone H3 (pH3, Blue). Scale bar: 50μm. (I) Model. Imp/Syp gradients regulate the metabolic switch through controlling the expression of mediator complex. Gradually eliminate Imp expression before the ecdysone signal (indicated by the open-box arrow) allows the metabolic switch that trigger non-MB neuroblast decommissioning. At last, Syp is required for the accumulation of Prospero to exit cell cycle.

Notably, Syp is required but not sufficient to increase Pros protein levels. For instance, a continuous induction of Syp since the beginning of neurogenesis failed to elicit precocious Pros accumulation in either neuroblasts or their progenies (data not shown). Co-induction of Imp and Syp throughout neurogenesis did not accelerate or delay Pros expression among serially derived progeny either, ascribing the increase of Pros in late-born neurons to other temporal cues independent of Imp/Syp. By contrast, ectopic Imp could effectively prevent high Syp from accumulating Pros in those long-lasting neuroblasts (Fig. 6C,D,H), arguing that nuclear accumulation of Pros depends on prior activation of neuroblast decommissioning rather than just high Syp. Considering all our data, we suggest that Syp plays a permissive role in the enhancement of Pros during late larval and early pupal neurogenesis.

## Discussion

Decommissioning of neuroblasts is a temporally controlled developmental program that proceeds from neuroblast shrinking to cell cycle exit. Knoblich’s group has shown that the mediator complex acts around pupation to uncouple the cell cycle from cell growth (Homem et al., 2014). This accompanies a change in cellular metabolism, which shrinks all but MB neuroblasts. It is also known that a subsequent nuclear accumulation of Pros protein results in neuroblast termination via a symmetrical division into two post-mitotic cells (Maurange et al., 2008). Here we show that neuroblast shrinking and cell cycle exit can be dissociated. In particular, knocking down Med12 to enhance mediator complex function causes Syp-depleted neuroblasts to shrink but not end (Fig. 4L). The long-lasting tiny neuroblasts could proliferate even in adult brains (Fig. 4L’’’’). This phenomenon indicates that the mediator complex is selectively involved in neuroblast shrinkage, and that neuroblast shrinkage does not automatically lead to cell cycle exit.

Imp can suppress neuroblast shrinkage and in turn prevent cell cycle exit (Figs. 1C,2A,3). By contrast, Syp is selectively required for neuroblast cell cycle exit. We established their respective roles by directly controlling both Imp and Syp to mitigate the fact that they negatively regulate each other (Liu et al., 2015). Silencing Imp made the ‘forever-young’ Syp-depleted neuroblasts shrink rapidly in early pupae but only moderately reduced the number of neuroblasts aberrantly present in adult brains (Fig. 1D). This ascribes the suppression of neuroblast shrinkage in Syp-depleted neuroblasts (Fig. 1B) to an increase in Imp rather than reduced Syp (Fig. 1F). It further demonstrates the requirement of Syp for neuroblast cell cycle exit. Interestingly, ectopic Imp can override the Syp-dependent cell cycle exit, as overexpressing Imp not only blocked neuroblast shrinkage but also prevented cell cycle exit even in the presence of enhanced Syp (Figs. 1C,G,6H). The dominant role of Imp effectively explains why MB neuroblasts that have prolonged Imp expression can escape the prepupal ecdysone pulse’s global initiation of neuroblast decommissioning. Consistent with this idea, repressing Imp, but not overexpressing Syp, made MB neuroblasts shrink rapidly in early pupae (Fig. 2). The MB neuroblasts with reduced cell size showed no evidence of cycling but could survive into late pupal stage despite high Syp levels. This might be due to the fact that MB neuroblasts end by apoptosis (rather than symmetric cell division) prior to eclosion (Siegrist et al., 2010).

A late larval induction of transgenic Imp could protect the normally ‘aged’ neuroblasts from decommissioning in early pupae. However, a slightly later induction around pupation failed to show any protection (Fig. 3). This assigns the critical window of Imp’s action to be around the prepupal ecdysone pulse. It suggests that the Imp itself (rather than the accumulative effects of protracted Imp expression) suppresses the initiation of neuroblast decommissioning by ecdysone. It further implies that neuroblasts do not age chronically due to repeated cycling.

Knocking down Med12 to enhance the mediator complex’s functions could largely erase the negative effects of ectopic Imp on neuroblast shrinkage and cell cycle exit (Fig. 4M). However, overexpressing Med12 alone only partially blocked the rapid shrinkage of the Imp-depleted MB neuroblasts (Fig. 4I-K). This argues that the prolonged Imp expression protects MB neuroblasts from early pupal decommissioning not only through enhancing Med12 but also via regulating other effectors to suppress the mediator complex. Consistent with this notion, we found that the *Med6* mRNA can be greatly enriched by Imp IP, implicating Med6 as an Imp direct target (Fig. 5). By contrast, Imp IP failed to pull down *Med12* mRNA, suggesting an indirect positive regulation of Med12 by Imp.

Besides, high Imp blocked neuroblast decommissioning to a much larger degree than inhibiting the mediator complex, suggesting involvement of mediator complex-independent targets. Taken together, the Imp RNA-binding protein may inhibit the mediator complex and other ecdysone effectors via multiple direct and indirect mechanisms to prevent neuroblasts from decommissioning until the production of almost all progeny.

Aged neuroblasts exit cell cycles due to nuclear accumulation of Pros. Drastic enhancement of Pros also occurs among the newborn cells from late larval and early pupal neuroblasts. Syp is required for the nuclear accumulation of Pros in aged neuroblasts as well as the strong Pros expression in late-born neurons (Fig. 6). However, high Syp alone is not sufficient to increase Pros protein levels, even though Syp protein may bind directly with Pros mRNA to enhance its stability and/or translation (McDermott et al., 2014). These observations argue that Pros is regulated at both transcriptional and posttranscriptional levels. As to timely cell cycle exit, a Syp-independent mechanism may upregulate Pros transcription at certain times following the activation of neuroblast decommissioning. High Syp would allow efficient translation of the increased Pros transcripts to flood the shrunk neuroblasts with copious Pros proteins, leading to nuclear accumulation of Pros and termination of the stem cell mode of asymmetric cell division.

Imp/Syp gradients in post-mitotic cells determine their birth order-dependent cell fates. Imp/Syp gradients in progenitors govern their readiness for aging and cell cycle exit. This allows decommissioning to be selectively activated in those neuroblasts that have produced essential progeny and are ready for aging. The opposing Imp/Syp temporal gradients with distinct lineage-specific temporal dynamics can therefore tailor neurogenic programs characteristic of different neuronal lineages (Fig. 6I). Notably, various stem cells across diverse species express analogous descending Imp gradients (Nishino, et al., 2013; Toledano, et al., 2012). It is possible that the mechanisms of progeny temporal fating and progenitor aging co-evolved on the pre-existing Imp gradients in neural stem cells. The achieved co-regulation of progeny temporal fate and progenitor decommissioning facilitates deriving complex protracted neuronal lineages.

## Material and Method

### Fly Strains and DNA constructs

The fly strains used in this study include (1) dpnEE-Gal4 [Awasaki et al., 2014; Yang et al., 2016]; (2) UAS-Syp-RNAi [stock #33011 and 33012, VDRC stock center];(3) UAS-Imp-RNAi [stock #34977 and 55645, Bloomington stock center]; (4) UAS-Imp-RM-Flag [Liu, et al., 2015]; (5) UAS-Syp-RB-HA [Liu, et al., 2015]; (6) UAS-Med12-RNAi [stock #34588, Bloomington stock center]; (7) UAS-pros.S [stock#32245, Bloomington stock center]; (8) lexAop-Syp-miRNA [Ren et al., submitted]; (9) UAS-Kto.J [stock #63801, Bloomington stock center]; (10) FRTG13, UAS-mCD8::GFP; GAL4-OK107 [Lee et al., 1999]. (11) Imp[CB04573] [gift from D.L. Jones, The Salk Institute for Biological Studies; Toledano, et al., 2012]. The following strains are generated in this study, (12) UAS-GFP::Sec; (13) dpn-LexA::P65; (14) act5C-GAL80ts-insulated spacer-tub-GAL80ts.

UAS-GFP::sec: the GFP fragment was inserted in a NotI/XbaI site in 5xUAS-pMUH plasmid (Awasaki et al., 2014), then the N-terminal 1-50 amino acids of securin was inserted in-frame at the C-terminus of GFP in an EcoRI/XhoI site. Dpn-LexA::P65: a previously characterized *dpn* neuroblast enhancer (Emery and Bier, 1995) and a *Drosophila* synthetic core promoter (Pfeiffer et al., 2008), was inserted in front of LexA::P65 in pBPLexA::P65w (Pfeiffer et al., 2010) through gateway cloning (Invitrogen).

act5C-GAL80ts-insulated spacer-tub-GAL80ts: a pGAL80ts plasmid was created by removing the HindIII/BglII fragment in pJFRC-MUH (Pfeiffer et al., 2010) then inserting the GLA80ts fragment from pMK-RQ(KanR)-GAL80ts into a KpnI/XbaI site. The *tubulin* promoter was inserted in front of GAL80ts in *pGAL80ts* through gateway cloning. The *actin5C* promoter was inserted in front of GAL80ts in *pGAL80ts* through gateway cloning. A synthetic insulated spacer cassette (Pfeiffer et al., 2010) was inserted at a FseI site in *pAct5C-GAL80ts*, and the EcoRI fragment of *pTub-GAL80ts* was then inserted in the EcoRI site after the insulated spacer cassette.

### Temporal induction of Pros

Flies with genotype UAS-pros.S; act5C-GAL80ts-insulated spacer-tub-GAL80ts/dpn-LexA::P65; 13XlexAop-Syp-miRNA/ dpnEE-Gal4 were cultured at 18°C. The GAL80 and temperature non-sensitive LexA::p65 allowed Syp knockdown continuously. For the control experiment, collected white pupae were cultured at 18°C for 24h before dissection. To induce Pros expression in neuroblasts, collected white pupae were transferred to 29°C to inactivate the temperature sensitive GAL80 for 16h before dissection, which resulted in the activation of Gal4 to express Pros.

### Temporal induction of Imp

Embryos with genotype act5C-GAL80ts-insulated spacer-tub-GAL80ts/dpn-Gal4; UAS-Imp-RM-Flag/UAS-GFP::Sec were collected for 24h at 18°C and then cultured at 18°C for 5 days. The larvae were heat shocked at 37°C for 15 min and incubated at 29°C to inactivate the temperature sensitive GAL80. White pupae were then collected at several time points (0h, 8h, 12h, 15h, 18h, and 24h) after heat shock, which resulted in the induction of Imp expression at different developmental time points (0h, 8h, 12h, 15h, 18h, and 24h) before pupal formation. Collected white pupae were cultured at 29°C and dissected at 38h APF.

### Immunohistochemistry and confocal imaging

Brain tissues at specific developmental stages were dissected and immunostained as described previously (Lin et al., 2012). The following primary antibodies were used: chicken anti-GFP 1:1000 (A10262, Life Technology), rat anti-Mira 1:200 (ab197788, abcam), rat anti-dpn 1:200 (ab195172, abcam), rabbit anti-pH3 1:500 (#9701, Cell Signaling), mouse anti-Pros 1:200 (Developmental Studies Hybridoma Bank), Guinea pig anti-Med12 1:500 (gift from J. Treisman, NYU school of medicine; Janody, et al., 2003), rabbit anti-Imp 1:600 (gift from P. Macdonald, University of Texas at Austin; Geng and Macdonald, 2006), and guinea pig anti-Syp 1:500 (McDermott et al., 2012). All corresponding fluorescent secondary antibodies (1:500) were purchased from Life Technology and images were collected on a Zeiss LSM 710 confocal microscope.

### Immunoprecipitation

Third instar larval brains dissected in Schneider medium were homogenized in 150 μl immunoprecipitation (IP) buffer (50 mM Tris-HCl pH 8.0, 150 mM NaCl, 0.5% NP-40, 10% glycerol, Complete EDTA-free protease inhibitor and RNAse inhibitor (RNAsin(®) Plus RNase Inhibitor (Promega)). The lysate was precleared with 45 μl of washed Pierce Control Agarose Resin (Thermo). For each reaction, 50 μl of pre-cleared lysate was taken as a 50% input sample directly to RNA extraction. 100 μl of pre-cleared lysate was incubated with 25 μl of GFP-TRAP agarose beads (Chromotek) for two hours at 4°C with rotation. The beads were washed 4 times briefly each with 200μl cold IP buffer at 4°C. After the final wash, beads were resuspended in 100 μl extraction buffer (50 mM Tris-HCl pH 8.0, 10 mM EDTA and 1.3% SDS, 1:100 RNAsin) and incubated at 65°C, 1000 rpm for 30 min in a thermomixer. The elution step was repeated and the supernatants were pooled. RNA was extracted from inputs and immunoprecipitates with the RNAspin RNA Isolation kit (GE Healthcare) and eluted in 40 μl of nuclease free H2O. Reverse transcription was performed using RevertAid Premium Reverse Transcriptase (Thermo Scientific) with random hexamer primers. cDNA was then used as a template for real time quantitative PCR.

### Real Time Quantitative PCR (RT-qPCR)

RT-qPCR was performed with SYBRGreen Mastermix (ThermoFisher) using a real time PCR detection system (CFX96 TouchTM Real-Time PCR Detection System (BioRad)). Cycle threshold (C(T)) value was calculated by the BioRad CFX software using a second differential maximum method. A dilution series of the input sample allowed the percentage input of each gene to be calculated to assess immunoprecipitation efficiency. The forward and reverse qPCR primer sequences are list in Table 1.

**Table 1.**
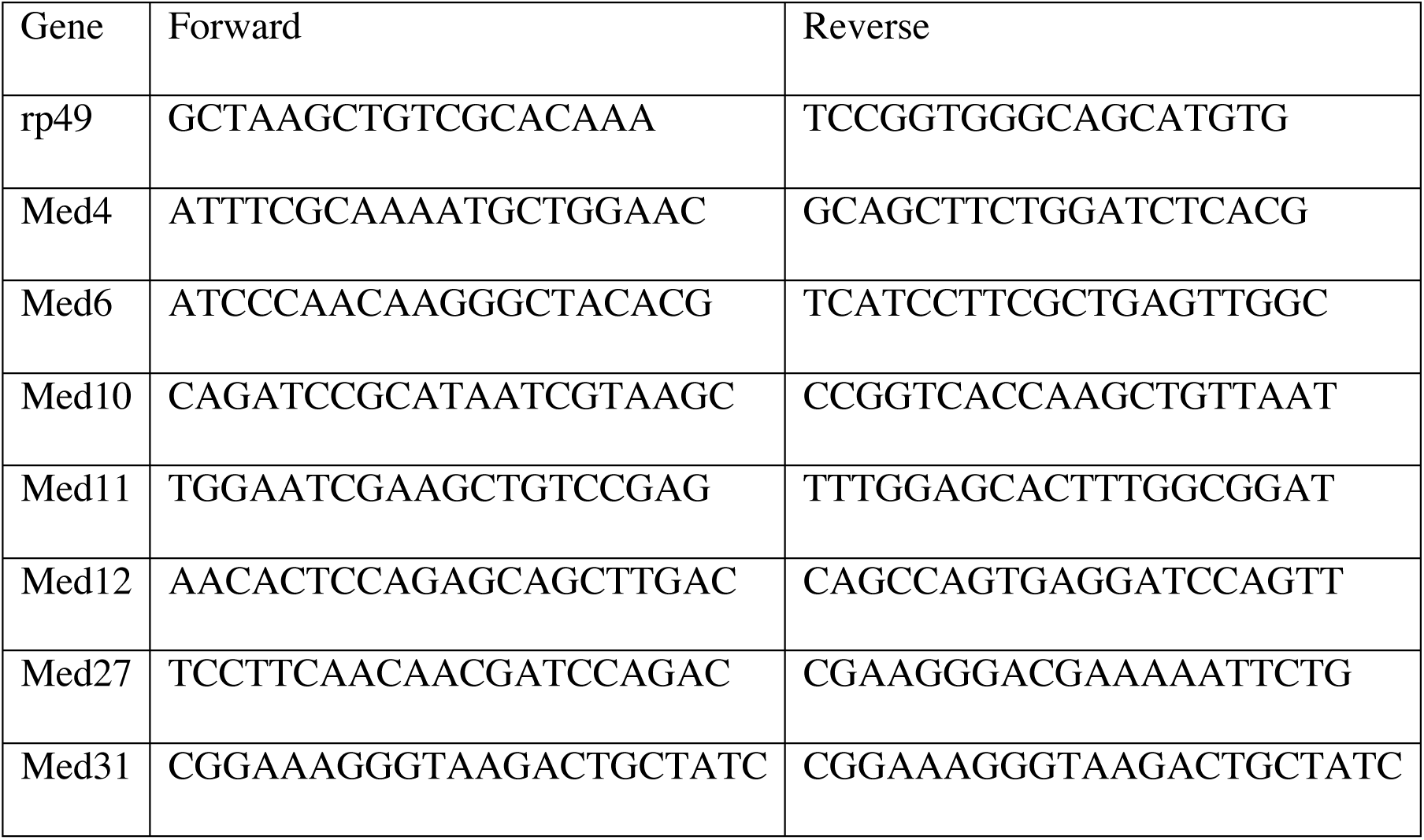
List of qPCR primer sequences.

## Acknowledgements

We thank the Janelia Fly Core, the Bloomington *Drosophila* stock center, the Vienna *Drosophila* Resource Center (VDRC) for technical support; J. Treisman for Med12 antibodies; P. MacDonald for Imp antibody; D.L. Jones for Imp[CB04573] line; R. Miyares for input and critical reading of the manuscript; J. Truman and D. Stern for helpful discussions and C. Di Pietro for administrative support.

### Competing interests

The authors declare no competing and financial interests.

### Author contributions

C.-P.Y. and T.L. conceived and designed the study. C.-P.Y. and T.J.S performed experiments. Y.H. made various DNA constructs. L.Y., D.I.H. and I.D. T.J.S. advised on experimental design and manuscript revision. C.-P.Y., T.J.S. and T.L. analyzed the data and C.-P.Y. and T.L. wrote the manuscript.

### Funding

C.-P.Y., Y.H. and T. L. were supported by the Howard Hughes Medical Institute. T.J.S. was funded by Wellcome Trust DPhil studentship (105363/Z/14/Z). L.Y. was supported by the Clarendon Trust and Goodger Fund. D.I.H. was funded by University College London. I.D. was funded from a Wellcome Senior Research Fellowship (081858) and the MICRON Oxford Wellcome Strategic Awards (091911/B/10Z and 107457/Z/15/Z). Deposited in PMC for release after 6 months.

